# Glucose-lactose mixture feeds in industry-like conditions: a gene regulatory network analysis on the hyperproducing *Trichoderma reesei* strain Rut-C30

**DOI:** 10.1101/2020.10.02.324319

**Authors:** Aurélie Pirayre, Laurent Duval, Corinne Blugeon, Cyril Firmo, Sandrine Perrin, Etienne Jourdier, Antoine Margeot, Frédérique Bidard

**Affiliations:** IFP Energies nouvelles, 1 et 4 avenue de Bois-Préau 92852 Rueil-Malmaison, France; ESIEE Paris, Université Paris-Est, Laboratoire d’Informatique Gaspard Monge (LIGM), 93162 Noisy-le-Grand, France; Genomic facility, Institut de Biologie de l’ENS (IBENS), Département de biologie, École normale supérieure, CNRS, INSERM, Université PSL, 75005 Paris, France

**Keywords:** *Trichoderma reesei*, carbon sources, cellulases, transcriptome, fed-batch fermentation, data science, Gene Regulatory Network

## Abstract

**Background:** The degradation of cellulose and hemicellulose molecules into simpler sugars such as glucose is part of the second generation biofuel production process. Hydrolysis of lignocellulosic substrates is usually performed by enzymes produced and secreted by the fungus *Trichoderma reesei*. Studies identifying transcription factors involved in the regulation of cellulase production have been conducted but no overview of the whole regulation network is available. A transcriptomic approach with mixtures of glucose and lactose, used as a substrate for cellulase induction, was used to help us decipher missing parts in the network.

**Results:** Experimental results confirmed the impact of sugar mixture on the enzymatic cocktail composition. The transcriptomic study shows a temporal regulation of the main transcription factors and a lactose concentration impact on the transcriptional profile. A gene regulatory network (GRN) built using the BRANE Cut software reveals three sub-networks related to *i* a positive correlation between lactose concentration and cellulase production, *ii* a particular dependence of the lactose onto the *β*-glucosidase regulation and *iii* a negative regulation of the development process and growth.

**Conclusions:** This work is the first investigating a transcriptomic study regarding the effects of pure and mixed carbon sources in a fed-batch mode. Our study expose a co-orchestration of *xyr1*, *clr2* and *ace3* for cellulase and hemicellulase induction and production, a fine regulation of the *β*-glucosidase and a decrease of growth in favor of cellulase production. These conclusions provide us with potential targets for further genetic engineering leading to better cellulase-producing strains.

## Background

Given current pressing environmental issues, research around green chemistry and sustainable alternatives to petroleum is receiving increased attention. A promising substitute to fossil fuels resides in second generation bio-ethanol, an energy source produced through fermentation of lignocellulosic biomass. One of the key challenges for industrial bio-ethanol production is to improve the competitiveness of plant biomass hydrolysis into fermentable sugars, using cellulosic enzymes.

The filamentous fungus *Trichoderma reesei*, because of its high secretion capacity and cellulase production capability, is the most used microorganism for the industrial production of cellulolytic enzymes. The *T. reesei* QM6a strain, isolated from the Solomon Islands during the Second World War [1], was improved through a series of targeted mutagenesis experiments [2, 3, 4, 5]. Among the variety of mutant strains, Rut-C30 is actually known as the reference hyper-producer [6, 7], and its cellulase production is 15-20 times that of QM6a [8]. Comparison of genomes of the Rut-C30 strain and its ancestor QM6a brings to light the occurrence of numerous mutations including 269 SNPs, eight InDels, three chromosomal translocations, five large deletions and one inversion [9, 10, 11, 12, 13, 14]. Alas among them, only few mutations have been proved to be directly linked to the hyper-producer phenotype [15, 10], the most striking one being the truncation of the gene *cre1* [9]. CRE1 is the main regulator of catabolite repression which mediates the preferred assimilation of carbon sources of high nutritional value such as glucose over others [16]. The truncated form retaining the 96 first amino acids and results in a partial release of catabolite repression [9] and more surprisingly turns CRE1 into an activator [17].

In *T. reesei*, the expression of cellulases is regulated by a set of various transcription. Beside the carbon catabolite repressor CRE1, the most extensively studied is the positive regulator XYR1 which is needed to express most cellulase and hemi-cellulase genes [18, 19]. Other transcription factors involved in biomass utilization have been characterized: ACE1 [20], ACE2 [21], ACE3 [22], BGLR [15], HAP 2/3/5 complex [23], PAC1 [24], PMH20, PMH25, PMH29 [22], XPP1 [25], RCE1 [26], VE1 [27], MAT1-2-1 [28], VIB1 [29, 30], RXE1/BRLA [31] and ARA1 [32]. Moreover, transcription factors involved in the regulation of cellulolytic enzymes have also been characterized in other filamentous fungi: CLR-1 and CLR-2 in *Neurospora crassa* [33] or AZF1 [34], PoxHMBB [35], PRO1, PoFLBC [36] and NSDD in *Penicillium oxalium* [37, 38]. Yet, their respective function has not yet been established in *T. reesei*. Among the mentioned regulators, some are specific to cellulases or xylanases genes, or to carbon sources while others are global regulators, *e.g.* PAC1, which is reported to be a pH response regulator. This profusion of transcription factors reveals the complexity of the regulatory network controlling cellulase production. Better understanding links between regulators could be a major key in improving the industrial production of enzymes.

Gene Regulatory Network (GRN) inference methods are computational approaches mainly based on gene expression data and data science to build representative graphs containing meaningful regulatory links between transcription factors and their targets. GRN may be useful to visualize sketches of regulatory relationships and to unveil meaningful information from high-throughput data [39]. We employed BRANE Cut [40], a Biologically-Related Apriori Network Enhancement method based on graph cuts, previously developed by our team. It has been proven to provide robust meaningful inference on real and synthetic datasets from [41, 42]. In complement to classical analysis, such as differential expression or gene clustering, the graph optimization of BRANE Cut on *T. reesei* RNA-seq is likely to cast a different light on relationships between transcription factors and targets.

While cellulose is the natural inducer of cellulase production, authors in [43] showed that, in *Trichoderma reesei*, the lactose is capable to play the role of cellulase inducer. For this reason, this carbon source is generally used in the industry to induce the cellulase production in *T. reesei*. Efficient enzymatic hydrolysis of cellulose requires the synergy of three main catalytic activities: cellobiohydrolase, endoglucanase and *β*-glucosidase. The cellobiohydrolases cleave D-glucose dimers from the ends of the cellulose chain. Endoglucanases randomly cut the cellulose chain providing new free cellulose ends which are the starting points for cellobiohydrolases to act upon, hydrolyze cellobiose to glucose, thereby preventing inhibition of the rest of enzymes by cellobiose [44]. It is well known that in *T. reesei*, *β*-glucosidase activity [45, 46] has generally been found to be quite low in cellulase preparations [47]. It causes cellobiose accumulation which in turn leads to cellobiohydrolase and endoglucanase inhibition. To overcome this low activity, different strategies have been experimented: supplementation of the enzymatic cocktail with exogenous *β*-glucosidase [48, 49], construction of recombinant strains overexpressing the native enzyme [50, 47, 51], expressing more active enzymes or modifying the inducing process to promote the production of *β*-glucosidase. This latest approach was performed by using various sugar mixtures to modify the composition of the enzymatic cocktail [52]. Thus, an increase of *β*-glucosidase activity in the cocktail can be achieved by using a glucose-lactose mixture, also favorable in terms of cost.

In the present study, fed-batch cultivation experiments of the *T. reesei* Rut-C30 strain, using lactose, glucose and mixtures of both were performed. As observed previously, productivity was increased with the proportion of lactose in the mixture and an higher *β*-glucosidase activity was measured in the mixture conditions compare to pure lactose. To explore the molecular mechanisms underlying these results, a transcriptomic study was performed at 24 h and 48 h after the onset of cellulase production triggered by the addition of the inducing carbon source lactose. An overall analysis reveals significant impact of lactose/glucose ratios on the number of differentially expressed genes and, to a lesser extent, of sampling times. According to the following clustering analysis, three main gene expression profiles were identified: genes up or down regulated according to lactose concentration and genes over-expressed in the presence of lactose but independently of its proportion in the sugar mix. Interestingly, expression profile of these genes sets overlaps productivity and *β*-glucosidase curve confirming a transcripomic basis of the pheno-types observed. As transcription factors were identified in all transcriptomic profiles, we decided to deepen our understanding on the regulation network operating during cellulase production in *T. reesei*. A system biology analysis with BRANE Cut network selection was carried out to inferred links between differentially regulated transcription factors and their targets. Results highlight three sets of subnetworks, one directly linked to cellulases genes, one matching with *β*-glucosidase expression and the last one connected to developmental genes.

## Results

### Cellulase production is increased with lactose proportion but *β*-glucosidase activity is higher in glucose-lactose mixture

In order to study its transcriptomic behavior on various carbon sources, *T. reesei* Rut-C30 was cultivated in fed-batch mode in a miniaturized experimental device called “fed-flask” [53], allowing us to obtain up to 6 biological replicates with minimal equipment. Cultures were first operated for 48 h in batch mode on glucose for initial biomass growth (resulting in around 7 g L^−1^ biomass dry weight), then fed with different lactose/glucose mixtures e.g. pure glucose (G_100_), pure lactose (L_100_), 75% glucose + 25% lactose mixture (G_75_-L_25_), and 90% glucose + 10% lactose mixture (G_90_-L_10_).

As expected, pure lactose feed resulted in highest protein production, with 2.6 g L^−1^ protein produced during fed-batch, at a specific protein production rate (q_P_) of 7.7 ± 1.1 mg g^−1^ h^−1^ (Figure 1A and 1B). Glucose feed resulted in almost no protein production (q_P_ 15 times lower than on lactose) but in biomass growth (4.2 g L^−1^ biomass produced during fed-batch, see Additional file 1) while glucose/lactose mixtures resulted in intermediate profiles, with 0.6 g L^−1^ protein produced on 10% lactose (G_90_-L_10_), and 1.4 g L^−1^ protein produced on 25% lactose (G_75_-L_25_). We then determined the filter paper and *β*-glucosidase activities at 48 h after the beginning of fed-batch (Figure 1C and 1D): filter paper activity is correlated to lactose amounts whereas *β*-glucosidase activity is higher in carbon mixture. The obtained results are in accordance with the ones obtained in [53], allowing us to assume the absence of residual sugar accumulation in the medium during the fed-batch.

**Figure 1.**
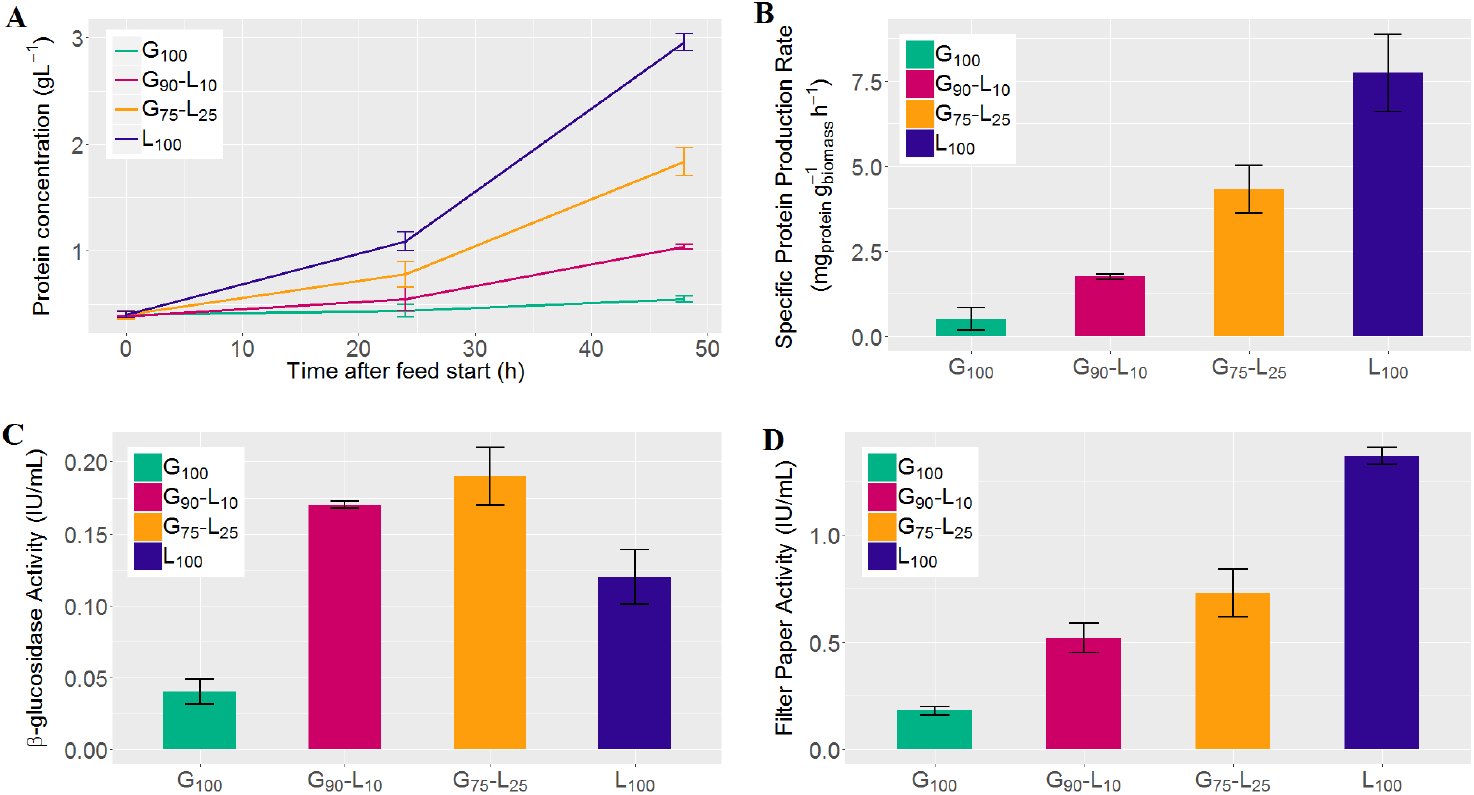
Protein production on different sugar sources in fed-batch mode. A: monitoring of protein concentration during fed-batch. For the different glucose-lactose content in feed (G_100_, G_90_-L_10_, G_75_-L_25_, L_100_), (B) reports the specific protein production rate, (C) the final *β*-glucosidase activity and (D) the final filter paper activity. Reported values are average and standard deviation of the biological replicates.

### Differentially expressed gene identification

This study aims at better understanding the effect of the lactose on the transcriptom of *T. reesei*, but not during the early lactose induction as in [54]. For this reason, we chose to extract RNA at 24 h and 48 h after the fed-batch start for further transcriptomic analysis.

Analysis of glucose, lactose and mixture effects was performed to identify differentially expressed (DE) genes between conditions. Specifically, to refine the understanding of the lactose effect on the cellulase production, the gene expressions on various lactose proportions (G_90_-L_10_, G_75_-L_25_, L_100_) at 24 h and 48 h have been differentially evaluated regarding gene expression obtained on pure sugar e.g. glucose (G_100_) or lactose (L_100_) at 24 h and 48 h. The comparison to both pure glucose and pure lactose feeds leads to ten comparisons (summarized on the circuit design displayed in Additional file 2. The use of two distinct references conditions increases the chances to identify relevant gene expression clusters by exploring a wider gene expression pattern. The number of DE genes obtained for each of the comparisons is displayed in Figure 2. For a better intelligibility of the results, we focus on DE genes compared to the pure glucose (G_100_) reference.

**Figure 2.**
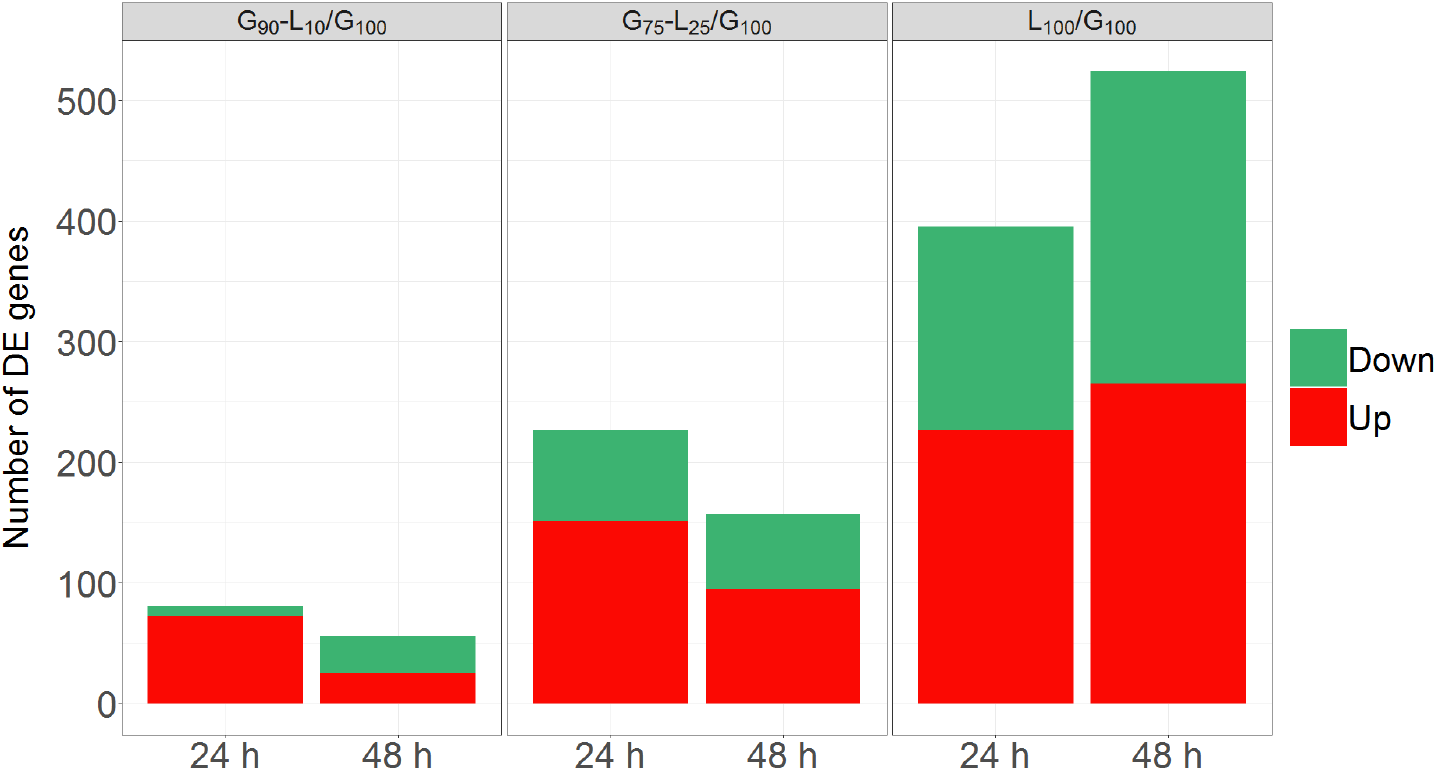
Differentially expressed genes of Rut-C30 on various of carbon sources mixtures. Number of over-(up, in red) and under-expressed (down, in green) genes on different mixed carbon source media (G_90_-L_10_, G_75_-L_25_, L_100_) at 24 h and 48 h.

From a global overview, at 24 h, 427 genes are differentially expressed and the number of DE genes increases with the level of lactose. In addition, these DE genes are up-regulated. Results obtained at 48 h lead to 552 DE genes and its number increases with the level of lactose. These results, displaying an increasing number of differentially expressed genes according to the lactose level between 24 h and 48 h, are in accordance with the specific protein production rate results previously presented (cf Figure 1). Note that this increase is essentially inherent to the threshold of 2 on the log fold-change. Indeed, at 24 h, some genes are considered as non differentially expressed although they are on the verge of becoming one, and then appear at 48 h.

We then focused on the intertwined effects *i.e.* the impact of time regarding each carbon source mixture. On pure lactose (L_100_), the number of DE genes increases between 24 h and 48 h. On the contrary, for both the minimal and the intermediate level of lactose (e.g. G_90_-L_10_ and G_75_-L_25_), the number of DE genes decreases between 24 h and 48 h. We observe that this diminution between the early and the late time samplings on low lactose quantity is mainly due to the diminution of over-expressed genes. This result suggests that a belated process only appears on pure lactose.

Eventually, we checked whether the genes mutated in Rut-C30, by comparison to QM6a, are differentially expressed in our conditions (see Additional file 3). While the total number of mutated genes at the genome scale is 166 (1.8%), we only found 12 of them in Rut-C30 which are also differentially expressed (1.8%). Hence, we cannot conclude to an enrichment of mutated genes responsible for cellulase production on lactose. This result is consistent with [54], which demonstrates the weak impact of random mutagenesis on transcription profiles related to cellulase induction and the protein production system.

Subsequent analyses are based on the 650 genes identified as DE in at least one of the ten studied comparisons.

### Gene clustering and functional analysis

To detect functional changes on lactose, we performed a clustering on the previously selected 650 genes. For this purpose, each gene is related to a ten-point expression profile corresponding to the ten log_2_ expression ratios (base-2 logarithm of expression ratios between two conditions according to the circuit design detailed in Additional file 2. Gene clustering was performed using an aggregated *K*-means classifier (detailed in the Materials and Methods section). Among the five distinct profiles identified (Figure 3 and Additional file 3 for the exhaustive list of genes), three main trends appear, when we compare the gene expression on lactose relatively to on glucose. The first trend encompasses genes under-expressed on lactose, in a monotonic manner at 24 h and 48 h and is found in two clusters, denoted by 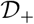 and 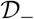 (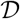 for down-regulation). Conversely, observed in two others clusters named 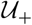 and 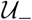 (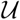 for up-regulation), the second trend refers to genes over-expressed on lactose in a monotonic manner at 24 h and 48 h. The last trend concerns genes over-expressed on lactose, but where the amount of lactose affects the gene expression in an uneven manner. This trend is recovered in a unique cluster denoted by 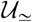.

**Figure 3.**
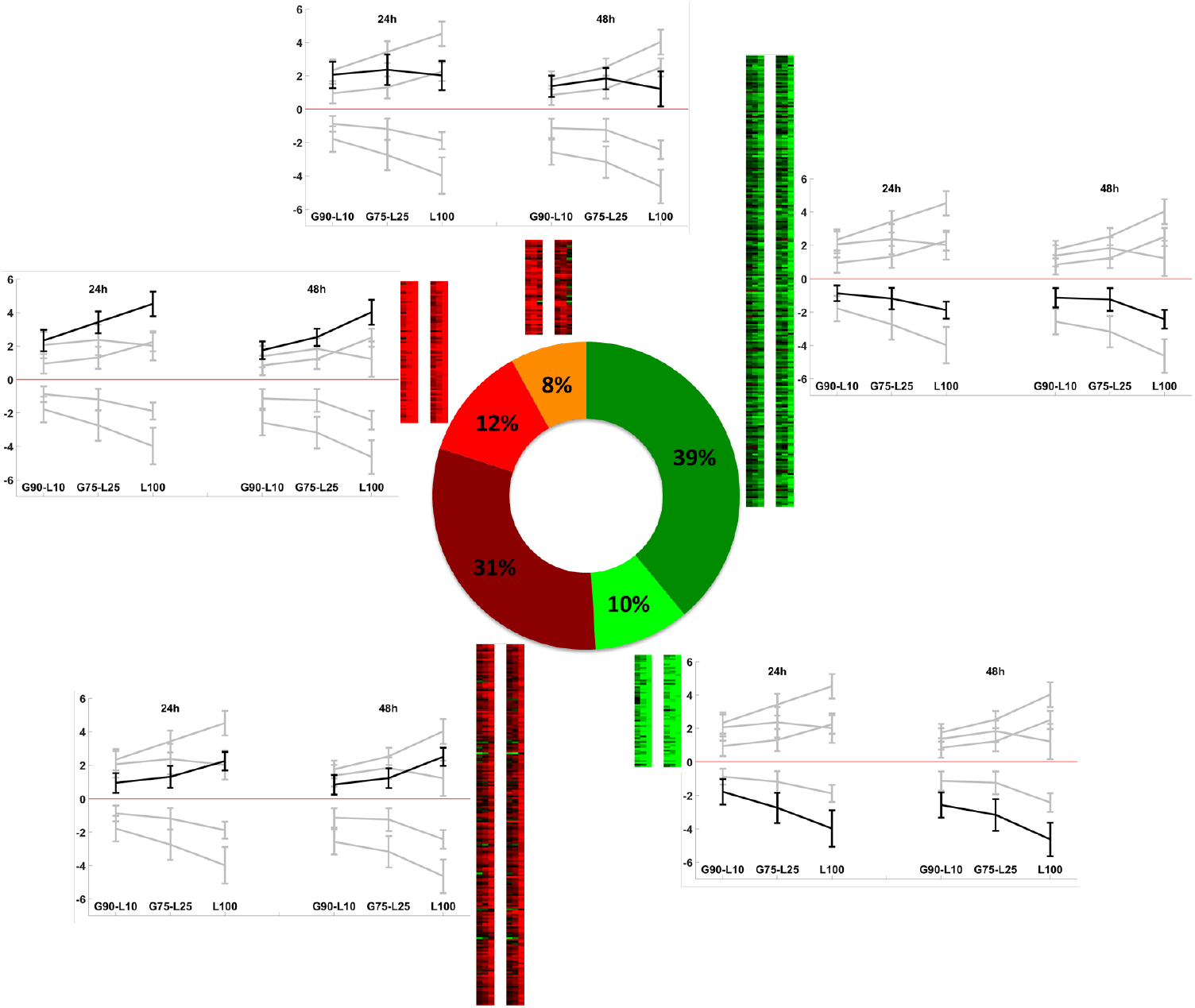
Heatmap and median profiles of clustered genes. Clustering results on the 650 differentially expressed genes : cluster 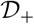 (green), 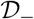 (dark green) for down-regulation, 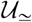 (orange), 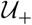 (red) and 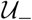 (dark red) for up-regulation. We have highlighted the median profile of the corresponding cluster in black and left the median profiles of the other clusters in grey in the background to facilitate visual comparison.

#### Genes monotonically down-regulated across lactose amount

As mentioned above, genes having a monotonic under-expression regarding the amount of lactose are grouped in clusters 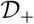 (64 genes: 10%) and 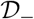 (254 genes: 39%). These genes are repressed in lactose: the more the lactose, the more the repression. The main difference between these two clusters is in the levels of under-expression: genes in cluster 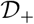 are in average more strongly under-expressed than genes in cluster 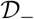. In addition, we note that cluster 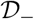, for which the under-expression is the weaker, contains a larger number of genes than cluster 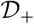. This result suggests that lactose moderately affects the behavior of a large number of genes while only few genes are strongly impacted by lactose concentration. In addition, it is interesting to note that the differential expressions of transcription factors are lower than genes not identified as such. This observation confirms that a weak modification only of transcription factors expression can lead to a strong modification in the expression of their targets.

More specifically, cluster 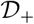 is enriched in genes related to proteolysis and peptidolysis processes (IDs 22210, 22459, 23171, 106661, 124051) and contains three genes encoding cell wall proteins (IDs 74282, 103458, 122127). Interestingly, no transcription factors are detected in this cluster.

Cluster 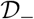, whose median profile exhibits a slight repression across lactose concentrations encompasses transcription factors whose ortholog are involved in the development: *Tr* –WET-1 (ID 4430, [55]), *Tr* –PRO1 (ID 76590, [56, 57]) and *Tr* –ACON-3 (ID 123713, [58]). We recall that the *Tr* –XXX notation refers to the gene in *T. reesei* for which the ortholog in an other specie is XXX (see the *Functional analysis* section in Materials and Methods). We also found 11 genes involved in proteolysis and peptidolysis processes, five genes encoding for cell wall protein (IDs 80340, 120823, 121251, 121818 and 123659), two genes encoding for hydrophobin proteins (*hbf2* and *hbf3*) and two genes involved in the cell adhesion process (IDs 65522 and 70021). Nine genes encoding for G-protein coupled receptor (GPCR) signaling pathway are also recovered in this cluster. It is important to note that, in addition to the three already mentioned, 11 other transcription factors are also present (including PMH29, RES1 [59], *Tr* –AZF-1 (ID 103275) and IDs 55272, 59740, 60565, 63563, 104061, 105520, 106654, 112085). We also found the xylanase XYN2 with a strong repression observed on pure lactose in comparison to pure glucose, while its expression seems insensitive to low lactose concentration.

#### Genes monotonically up-regulated across lactose amount

We recall that clusters 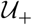 (78 genes: 12%) and 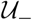 (201 genes: 31%) contain genes whose over-expression is monotonic with respect to lactose: the more the lactose, the more the induction. The main difference between expression profiles of these two clusters is the level of over-expression: genes in cluster 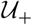 are more activated than genes belonging to cluster 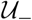. A similar remark may be drawn as previously: preliminary observations suggest that a large number of genes is moderately impacted by lactose (cluster 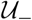) while only few genes are strongly affected by lactose concentrations (cluster 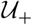). As similarly observed on down-regulated genes, the expression level of the transcription factors is weaker than their targets.

In cluster 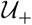, whose median profile expresses a potent induction regarding lactose concentrations, 26 CAZymes are found, of which 23 belong to the large glycoside hydrolase (GH) family. We recover the principal CAZymes known to be induced in lactose condition: the two cellobiohydrolases CBH1 and CBH2, two endoglucanases CEL5A and CEL7B, one lytic polysaccharide monooxygenase (LPMO) CEL61A, two xylanases XYN1 and XYN3, as well as the mannanase MAN1, the *β*-galactosidase BGA1. In addition, we found three specific carbohydrate transporters CRT1, XLT1 and ID 69957 and three putative ones (IDs 56684, 67541, and 106556). Interestingly, we found the transcription factor YPR1, which is the main regulator for yellow pigment synthesis [60]. These results, showing a lactose-dependent increase in the expression of genes related to the endoglucanase and cellobiohydrolase, corroborate the phenotype observed in the study of [52]. Indeed, its authors show a rise of the specific endoglucanase and cellobiohydrolase activity positively correlated to lactose concentration and cellulolytic enzymes productivity.

Cluster 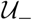, distinguishable by its median profile showing a slight induction across lactose concentrations, contains 17 genes involved in the carbohydrate metabolism, of which 16 belong to the large GH family. Among these genes, we identified three *β*-glucosidases whose two extracellulars CEL3D and CEL3C and one intracellular CEL1A, the xylanase XYN4, and the acetyl xylanase esterase AXE1 are recovered. We also found 14 Major Facilitator Superfamily (MFS) transporters. In addition, seven transcription factors are found in this cluster, including XYR1 the main regulator of cellulase and hemicellulase genes [19], CLR2 (ID 23163) identified as a regulators of cellulases but not hemicellulases in *Neurospora crassa* [33], *Tr* –FSD-1 (ID 28781), ID 121121 and three others, with no associated mechanism (IDs 72780, 73792, 106706).

#### Uneven up-regulation across lactose amount

In cluster 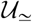 (53 genes: 8%), we found globally over-expressed genes but with a non-monotonic behavior regarding lactose concentration. A more detailed study of this cluster reveals three main typical characteristics in the gene expression profiles. A tenth of the genes shows an uneven behavior with a high-over expression in all G_90_-L_10_, G_75_-L_25_ and L_100_ conditions without significant difference according to the amount of lactose. This kind of profile suggests that the up-regulation is uncorrelated with lactose concentration itself but triggered by lactose detection only. Then we found one third of the genes that demonstrates a high over-expression on the two carbon source mixtures G_90_-L_10_ and G_75_-L_25_ while no differential expression is observed on pure lactose compared to pure glucose. The transcription factor ID 105805 follows this profile. Finally, a little more than half of the genes has a significant stronger over-expression on G_75_-L_25_ compared to the one on G_90_-L_10_ and L_100_. Interestingly, we found one endoglucanase CEL12A, one LPMO CEL61B, three *β*-glucosidases whose two extracellulars with a peptide signal CEL3E and BGL1 and one intracellular *β*-glucosidase CEL1B, potentially involved in cellulase induction. We also found the *β*-xylosidase BXL1 and the transcription factor ACE3 that share this profile. We observe a strong correlation between the transcriptomic behavior we found in our study and the phenotype highlighted in [52]. Actually, the specific *β*-glucosidase activity is the highest for intermediate amounts of lactose while this activity decreases on glucose or lactose alone. Corroboratively, our transcriptomic study shows a highest over-expression of genes encoding *β*-glucosidases (*cel3e*, *bgl1* and *cel1b*) on the intermediate mix of lactose and glucose, while their expression decreases when lactose is present in too low or too high concentration.

Note that a large proportion of genes belonging to the up-regulated clusters are recovered on the co-expressed genomic regions observed in [22]. The biological coherence of clustering results encourage us to pursue the transcriptomic study through a gene regulatory network. The use of network inference approach is driven by the motivation to better understand links between DE transcription factors but also to highlight strong links with the help of alternative proximity definition, and thus to concrete the relationships foreseen though the clustering.

### Network inference

From the set of DE genes, we built a gene regulatory network with the combination of CLR [61] and BRANE Cut [40, 62] inference methods. When the use was judicious, we evaluated our discovered TF-targets interactions by performing a promoter analysis of the plausible targets given by the inferred network, with the Regulatory Sequence Analysis Tool (RSAT) [63]. More details on the complete methodology for both the inference and the promoter analysis are provided in section Materials and Methods.

Network enhancement thresholding performed by BRANE Cut post-processing [40] selected 161 genes (including 15 transcription factors) and inferred 205 links (Figure 4). In order to help network interpretation, we applied the same color code as for the clustering (Fig. 3). We observe a coherence between the function and the expression behavior of genes linked into modules, thus corroborating clustering results. As we will see in details in the following network analysis, we reveal potential links between three mechanisms grouped in modules (SubN_1_, SubN_2_, and SubN_3_) and related to cellulase activation, *β*-glucosidase expression and repression of developmental process.

**Figure 4.**
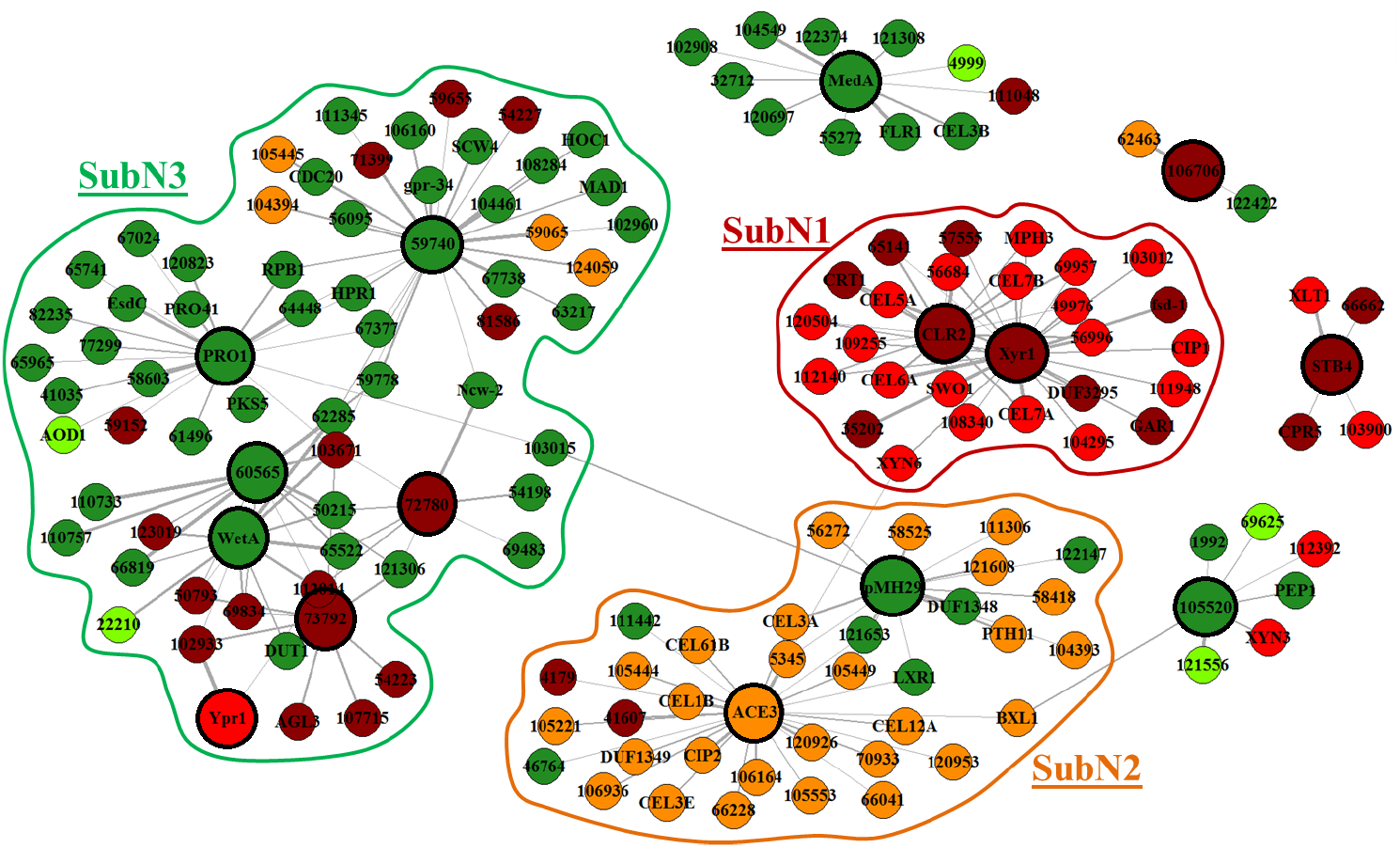
Inferred network. Network built with BRANE Cut from expression profiles of the differentially expressed genes. BRANE Cut selected 205 edges involving 161 genes. Node colors correspond to cluster labels: 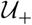 (red, genes highly and monotonically up-regulated on lactose), 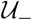 (dark red, genes slightly and monotonically up-regulated on lactose), 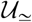 (orange, genes up-regulated and non-monotonically on lactose), 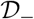 (dark green, genes slightly and monotonically down-regulated on lactose) and 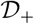 (green, genes highly and monotonically down-regulated on lactose). Bigger nodes with bold frame correspond to genes coding for a transcription factor while smaller nodes with thin frame correspond to genes not identified to code for a transcription factor.

First of all, the global study of the network shows interactions between genes sharing the same gene expression profile. The 161 genes selected by BRANE Cut cover a relatively small number of biological processes, especially regarding half of the 15 retained transcription factors for which only two main biological processes are identified: development (*Tr* –WET-1, *Tr* –PRO1, *Tr* –ACON-3 (IDs 4430, 76590, 123713)) and carbohydrate mechanisms (XYR1, PHM29, ACE3 and CLR2).

In addition, we observe a large proportion of genes related to the enzymatic cocktail for cellulase production. In terms of interaction, we predominantly observed links between up-regulated genes in a monotonic manner (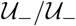 and 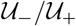 interactions), and related to cellulase production. A second observation refers to enriched 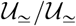 interactions i.e. between up-regulated genes in an uneven way. Note that we also found an interesting proximity with 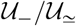 interactions, with inverse expression profiles. Involved genes mainly refer to the cellulase and *β*-glucosidase production. Finally, a significant number of interactions are found between genes belonging to cluster 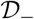 and related to development mechanism. Here again, links are also observed between genes having antagonist expression profiles, mainly related to cellulase production and development (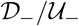 interactions). Figure 4 displays the inferred network with highlights on the three sub-networks SubN_1_, SubN_2_ and SubN_3_, extracted from the combination of the above observations and the clustering results. We now focus on each identified sub-network for a more detailed analysis.

Sub-network SubN_1_ encompasses eight genes associated to the carbohydrate metabolism process. Among them, *cel5a*, *cel6a*, *cel7a* and *cel7b* are specifically related to cellobiohydrolase and endoglucanase activities. It also includes four carbohydrate transporters including CRT1, responsible for lactose uptake, and three carriers [22, 64]. These genes are linked to transcription factor XYR1, known to be the main actor during the cellulase production process. It also appears specifically linked to a galacturonic acid reductase GAR1, a helicase (ID 35202), a glycoside hydrolase XYN6 [65], a secreted hydrolase CIP1 and *Tr* –FSD-1 (ID 28781), known to pertain to sexual development. The network highlights the action of another transcription factor CLR2, which is known in other species to participate to cellulase production [33]. These two transcription factors XYR1 and CLR2 seem to be highly correlated and share a large number of cellulose-oriented targets. This sub-network is related to the genes involved in cellulase production and having an increased up-regulation across to the lactose concentration. Based on this sub-network subN1, we performed a promoter analysis. Using independently plausible targets of XYR1 and CLR2, we significantly recovered the degenerated binding-site 5’-GGC(A/T)_3_-3’, previously identified in [66] as the binding site specific to XYR1. We also found an enriched non-degenerated motif 5’-GTTACA-3’ which differs from the XYR1 motif. A straightforward hypothesis is to credit this new motif for CLR2 and a simple statistical test suggests that this motif might be specific to the CLR2. Details regarding this analysis are provided in Additional file 4. To do.

Sub-network SubN_2_ contains nine genes involved in the carbohydrate metabolism, and some of them are specifically related to *β*-glucosidase and cellulases activities: *bgl1*, *cel3e*, *cel12a* and *cel61b*. Interestingly, these genes are linked to the transcription factor ACE3 and have the particularity to be maximally over-expressed on G_75_-L_25_. We observe that seven genes belonging to cluster 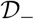 are also present in this sub-network and are predominantly linked to the transcription factor PHM29 which has been recently identified to play a role in the cellulase activity [22]. We notice that these genes have a maximal under-expression on G_75_-L_25_, which is the inverse profile of *ace3* and its linked genes, suggesting a dependence between ACE3 and the transcription factor *pmh29*.

The sub-network SubN_3_ reveals seven transcriptions factors including two which have been identified to participate to the development process in other species: *Tr* –WET-1 (ID 4430) and *Tr* –PRO1 (ID 76570). Interestingly, three other genes *EsdC*, *pro41* and *hpr1* , also pertaining to the development process, are linked to *pro1* . In addition, genes in this sub-network are mainly down-regulated on lactose and related to metabolism, secretion, transport and cell surface. This sub-network seems to reveal some interesting links between the repression of the development and the cellulase production that will be investigated in more details in the Discussion.

Results provided by this inferred network and the promoter analysis are in agreement with present knowledge on *Trichoderma reesei*, particularly for the cellulase production. The additional results given by BRANE Cut are coherent with the literature based on other close species, especially regarding results that suggest a potential link between development and cellulase production and a particular behavior of the *β*-glucosidase. Table 1 provides some relevant references that coroborate the network generated by BRANE Cut. The coherence of the DE analysis as well as clustering and inference results with the actual knowledge allows us to use these results for prediction. In the following Discussion section, we thus formulates some postulates regarding cellulase production mechanism in *T. reesei* , with respect to these three main results.

**Table 1.**
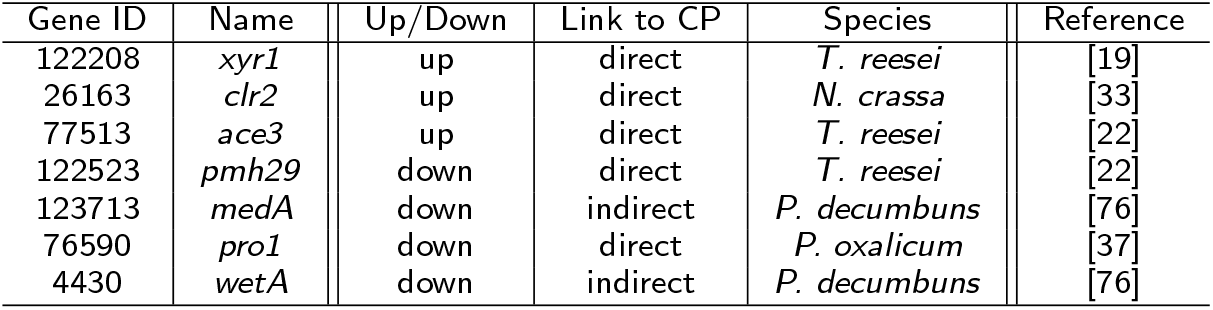
BRANE Cut network validation from literature. Direct link refers to genes identified as implied in the cellulase production while indirect refers to genes having a side effect on the cellulase production (CP).

| Gene ID | Name | Up/Down | Link to CP | Species | Reference |
| --- | --- | --- | --- | --- | --- |
| 122208 | *xyr1* | up | direct | *T. reesei* | [19] |
| 26163 | *clr2* | up | direct | *N. crassa* | [33] |
| 77513 | *ace3* | up | direct | *T. reesei* | [22] |
| 122523 | *pmh29* | down | direct | *T. reesei* | [22] |
| 123713 | *medA* | down | indirect | *P. decumbens* | [76] |
| 76590 | *pro1* | down | direct | *P. oxalicum* | [37] |
| 4430 | *wetA* | down | indirect | *P. decumbens* | [76] |

## Discussion

### A cellulase production directly linked to the lactose concentration

The gene *xyr1* is widely reported to play the role of the major activator of the cellulase production in *T. reesei* [19]. As notably expected, we recovered in our network links between XYR1 and the main cellulolytic enzymes (especially the two main cellulases CBH1 and CBH2). In *Neurospora crassa*, cellulases are regulated by CLR-2 specifically, while *Tr* –XLNR, the ortholog of *xyr1*, is responsible of the hemicellulase expression [33, 67]. Thus, the regulation of cellulases and hemicellulases is performed through two independents pathways. While the genes responsible for this regulation are present in *T. reesei* , their behavior appears to be different as they show a coupling action of the regulation of both cellulases and hemicelullases, suggesting a different regulatory network in *T. reesei* compared to *N. crassa*.

Although observed in different *T. reesei* strains and culture conditions, authors in [68] and [69] have identified links between *xyr1* and *clr2* genes. Interestingly, we also found in our data such a strong correlation between *xyr1* and *clr2*, suggesting a common regulation on lactose. We found a significant number of regulatory links between *clr2* and cellulolytic enzymes. Unlike in *N. crassa*, *clr2* seems to be complemental to *xyr1* for cellulases and hemicellulases activation in *T. reesei*. Thus, even though gene ID 26163 is the ortholog of *clr-2* in *N. crassa*, this observation argues for a different behavior in *T. reesei*.

Another difference between *T. reesei* and *N. crassa* regarding *clr2* is its location on the genome. Contrary to *N. crassa*, *clr2* in *T. reesei* pertains to a physical cluster, located on chromosome III [70], and containing the lactose permease CRT1, established as essential for cellulase induction on a lactose substrate as it allows lactose uptake [22, 64]. Due to this proximity between *clr2* and *crt1*, we may assume a regulation of *crt1* by CLR2. In *N. crassa*, the ortholog of *crt1* is *sud26* , and encodes a sugar transporter which is located next to a transcription factor of unknown function TF-48.

In *N. crassa*, *clr2* is repressed by the carbon catabolite repression [33]. We do not know if such an extrapolation to *T. reesei* is valid, but interestingly, the Rut-C30 strain has a partial release of catabolite repression due to the truncation of *cre1*, allowing us to suggest a possible release of the repression of *clr2* , leading to a basal expression of CLR2 and CRT1, so a basal lactose uptake. This low level of lactose would be sufficient to initiate the induction of cellulases through the expression of XYR1 and CLR2.

As established in [22], the gene *ace3* is known in *T. reesei* to be involved in the cellulase induction on lactose. Furthermore, as presented in [71], *ace3* seems to interact with *xyr1* to initiate cellulase production. Based on your data and their interpretations, especially regarding the strong correlation between *clr2* and *xyr1*, we may suppose an additional interaction between *ace3* and *clr2* . This result can also be corroborated by the fact that the invalidation of *ace3* in [71] leads to a decrease of XYR1 and CLR2 expressions. However, we note that the expression of ACE3 is not directly correlated with the lactose concentration as the maximal expression of ACE3 is obtained on a mixture of glucose and lactose (G_75_-L_25_). Thus, the regulation of XYR1 by *ace3* could be complemented by another mechanism necessary for cellulase induction on pure lactose, and without glucose.

### Gene expression profiles of *bgl1*, *cel3e* and *cel1b* follow *β*-glucosidase activity

A previous study had shown an effect of sugar mixtures to influence the composition of the enzymatic cocktail of *T. reesei* [52]. A higher *β*-glucosidase activity was observed in the presence of a glucose-lactose mixture compared to pure lactose. This result obtained in the CL847 strain is here confirmed in the reference hyper-producing Rut-C30 strain.

In the transcriptome performed on the various glucose-lactose mixtures, a group of DE genes (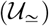) has an expression profile correlated to *β*-glucosidase activity, i.e. genes overexpressed by lactose but without correlation with the amount of lactose. Among these genes, three *β*-glucosidase are identified, whose two are extracellular (*bgl1* and *cel3e*) while the other is an intracellular *β*-glucosidase (*cel1b*). It has been shown previously that in presence of lactose the extracellular enzyme activity is mainly produced by *bgl1* [72]. Our results seem to demonstrate that for a full expression of *bgl1*, presence of lactose is required independently of glucose. Nothing is known about the regulation of *cel3e* but its expression profile is similar to *bgl1* . This two genes have been previously identified as co-regulated by the same substrate [73]. There is therefore a correlation between the expression of these genes and enzymatic activity of BGL1. It would thus be interesting to delete *cel3e* to study the impact of its absence on the global extracellular *β*-glucosidase activity in glucose-lactose mixture.

In the regulatory network, *bgl1* and *cel3e* are connected to both *ace3* and *pmh29* . However, *ace3* has a similar profile as the previously mentioned *β*-glucosidase (*cel3a* and *cel3e*) while *pmh29* is anti-correlated. It would therefore be interesting to explore the role of its two transcription factors in the control of CEL3A/BGL1 and CEL3E under glucose-lactose induction conditions. The roles of *ace3* and *pmh29* in cellulase regulation have recently been explored [22]. However, the difference in genetic background (QM6a and QM9414) and experimental conditions (100% lactose batch) does not allow the results of these experiments to be extrapolated to the regulation observed here.

Another *β*-glucosidase, CEL1B, is present in cluster 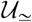. This intracellular enzyme appears to play an essential role in lactose induction since the joint invalidation of *cel1b* and *cel1a*, another intracellular *β*-glucosidase, abolishes the production of cellulases on lactose. However, invalidation of *cel1b* alone does not appear to have any effect while invalidation of *cel1a* produces a delay in induction on lactose which is restored by galactose. Surprisingly, the transcriptomic profile of *cel1a* is different from that of *cel1b* since it belongs to the cellulase cluster 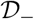. The difference in its profiles could indicate a different response between these two genes depending on whether or not glucose is present. Thus the expression of CEL1A could be negatively regulated by the presence of glucose and induced by lactose while CEL1B could be induced by lactose but insensitive to the presence of glucose. As *cel1b* is also connected to the regulators ACE3, it would be interesting to explore the role of ACE3 and PMH29 regulators in the expression of CEL1B.

### A dedication to cellulase production to the detriment of growth

Strinkingly, orthologs of transcription factor genes (IDs 4430, 76590 and 123713) described as involved in developmental process have been identified in this transcriptomic study. All of them being part of cluster 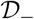 and so down-regulated in lactose compared to glucose.

Firstly, ID 76590 is the ortholog of *pro1* in *Sordaria macrospora* (67% identity) and *Podospora anserina* (49% identity), and the ortholog of *adv-1* in *Neurospora crassa* (67% identity). The gene *Tr* –*pro1* is required for fruiting body development and cell fusion [56, 57]. In *P. anserina*, *pro1* activates the sexual recognition pathway including the pheromone and receptor genes and is probably involved in the control of the entry in stationary phase [74]. In *Penicillium oxalicum*, deletion of *pro1* (43% identity) has been proved to increase cellulase production [37]. No similar phenotype has been described in other fungi. At low lactose concentration obtained in our experiments, *Tr* –*pro1* is down regulated and linked in the GRN to *hpr1*, the mating type pheromone receptor.

Secondly, ID 123913 is the ortholog *MedA* in *Aspegillus nidulans* (42% identity), coding for a protein with unknown function, but required for normal asexual and sexual development. We determined that the *N. crassa* ortholog of *MedA* is *acon-3*, a gene required for early conidiophore development and female fertility. In *N. crassa*, *acon-3* is positively regulated by the transcription factor ADA-6 involved in conidiation, sexual developement, and oxidative stress response [58]. Interestingly, *ypr1* (ID 102499), the yellow pigment regulator, DE in our data, displayed 35% identity with *ada-6* . In contrast to *Tr* –*MedA*, *ypr1* is up-regulated on lactose and its regulatory function seems restricted to the sorbicillin cluster [60].

The gene with ID 4430 is the ortholog of *wet-1* of *N. crassa* (72% identity), of *WetA* in *A. nidulans* (60% identity) and *Fusarium graminearum* (43% identity). In contrast to *Aspergilli* and *F. graminearum*, *wet-1* mutant is phenotipically similar to the wild type strain with no conidiation defect [55]. A regulatory cascade with *WetA* regulated by *AbaA* itself regulated by *BrlA* was described in *Aspergillus* [75]. In *P. decumbens*, an industrial lignocellulolytic enzymes production strain, expression of cellulases genes is upregulated in *BrlA* deletion strain [76]. In *T. reesei*, while no ortholog has been identified ofr *BrlA*, *rxe1* (20% identity with *BrlA*) is involved in regulation of conidiation and modulated positively by the expression of *xyr1* and cellulase and hemicellulase genes. The regulatory cascade between *aba1* and *wet-1* is preserved in *N. crassa* and *F. graminearum* but we do not know if the *rxe1* gene could replace *BrlA* in species where there is no true ortholog and therefore if *wet-1* may be controlled by *rxe1* . In our transcriptomic data, neither *rxe1* nor *Aba1* is differentially regulated, so down-regulation of *wet-1* does not seem to be dependent of these genes. Eventually, further experiments would allow us to decipher the role of *wet-1* on cellulase production and the regulatory link between *wet-1* and *rxe1* .

In *Aspergillus nidulans*, MEDA acts as a repressor of *BrlA* expression and is an activator of *AbaA* expression [77]. Although, no direct regulation relation between *MedA* and *WetA* in *T. reesei* has been described, it is worth to note that these genes, both involved in the regulation of conidiation, are down-regulated on lactose. Interestingly in *A. niger*, authors in [78] showed that the secretion of the vegetative mycellium is repressed by sporulation, thus indicating a reverse link between conidiation and secretion. Thus, *Tr* –*WetA* and *Tr* –*MedA* down-regulation could be a result of the lactose fed batch cultivation mode where the carbon flux is maintaining a near-vegetative state without growth. Conversely, glucose feed resulted in biomass growth leading to conidiation.

Altogether, the down regulation of *Tr* –*pro1*, *Tr* –*wet1* and *Tr* –*acon3* on lactose compared to glucose could reflect a balance between vegetative growth, sexual and asexual development. In the fed-batch condition, the lactose is provided to maintain the biomass without growth. In contrast, starvation could create a path to conidiation or glucose could redirect to sexual development. The equilibrium is maintained through the down regulation of essential developmental transcription factor.

## Conclusions

This study is the first considering the effect of various carbon sources (glucose/lactose mixtures) in a fed-batch mode on the *Trichoderma reesei* transcriptome. In such a condition, we highlighted an interdependence between crucial transcription factors (XYR1, CLR2 and ACE3) known to participate to cellulase and hemicellulase production. We also correlated the transcriptome to the *β*-glucosidase activity observed in a previous study [52] and revealed a repression of the development process during the cellulase production. These conclusions provide us with plausible targets for further genetic engineering leading to better cellulase producing strains.

## Materials and Methods

### Strain and media

*T. reesei* RUT-C30 (ATCC 56765) was received from ATTC on October 2013, spread on PDA plates and incubated until sporulation. Spores were harvested with 50% glycerol solution then stored at −80 °C. Spore solution concentration was 6 × 10^9^ mL^−1^. Culture media are prepared according to [53] (case with 25 mM dipotassium phthalate) and supplemented with 12.5 g L^−1^ glucose. Feeding solutions (stoichiometric mix of carbon and nitrogen sources) were prepared according to [52].

### Fed-flask cultivations

Fed-flask cultivation was performed according to [52] with few modifications. For each replicate, a Fernbach flask was prepared with 250 mL culture medium and inoculated with around 10^7^ spores mL^−1^. Initial growth phase on glucose lasted around 48 h and resulted in around 7 g L^−1^ biomass. Immediately after glucose exhaustion, empty 250 mL Erlenmeyer flasks were filled with 50 mL broth per flask then fed at 0.3 mL h^−1^ (using Dasgip MP8 peristaltic pumps) with different sugar solutions (one flask fed with pure lactose, one flask fed with pure glucose, one flask fed with a mixture of glucose and lactose). Pure glucose (G_100_) feed and pure lactose (L_100_) feed were replicated 6 times, 75% glucose + 25% lactose mixture (G_75_-L_25_) was replicated 4 times, and 90% glucose + 10% lactose (G_90_-L_10_) was replicated 2 times. Incubation was performed in an Infors rotary shaker at 30 °C and 150 rpm. Analysis (biomass dry weight, protein concentration, sugars concentration, enzymatic activities) were performed according to [52].

### RNA-seq library preparation and analysis

#### Library preparation and RNA-seq data acquisition

Library preparation and Illumina sequencing were performed at the École normale supérieure Genomic Platform (Paris, France). Messenger (polyA+) RNAs were purified from 1 μg of total RNA using oligo(dT). Libraries were prepared using the strand specific RNA-seq library preparation TruSeq Stranded mRNA kit (Illumina). Libraries were multiplexed by 6×6 flowcell lanes. A 50 bp read sequencing was performed on a HiSeq 1500 device (Illumina). An average of 35 ± 10 millions passing Illumina quality filter reads was obtained for each of the 36 samples. The analysis were performed using the Eoulsan pipeline [79], including read filtering, mapping, alignment filtering, read quantification, normalisation and differential analysis. Before mapping, poly N read tails were trimmed, reads with less than 40 bases were removed, and reads with quality mean lower than 30 were discarded. Reads were then aligned against the *Trichoderma reesei* genome version 2 (from the Joint Genome Institute database) using Bowtie (version 0.12.9) [80]. Alignments from reads matching more than once on the reference genome were removed using samtools (Java version) [81]. To compute gene expression, *Trichoderma reesei* genome annotation version 2 from Joint Genome Institute database was used. All overlapping regions between alignments and referenced exons (or genes) were counted using HTSeq-count 0.5.3 [82]. The RNA-seq gene expression data and raw fastq files are available on the GEO repository (www.ncbi.nlm.nih.gov/geo/) under accession number: GSE82287.

#### Normalization and differentially expressed genes identification

RNA-seq data normalization and differential analysis was performed thanks to the DESeq Bioconductor R package (version 1.8.3) [83]. The normalization method implemented in DESeq assumes that only a few number of genes are differentially expressed and corresponds to a median scale normalization.

The differential analysis relies on a statistical model, and more precisely on the negative binomial distribution with variance and mean related by local regression. This approach allows us to identify, for each gene, if the observed difference in read counts is significant. An adjustment for multiple-testing with the procedure of Benjamini and Hochberg [84] was also performed. Hence, we assumed that a gene is said differentially expressed when the adjusted *p*-value was lower than 0.001 and the absolute value of the log_2_pFCq was higher than 2. Here, FC refers to the fold change of the read counts for the tested condition against the read counts for the reference condition. In this way, we independently compared at 24 h and 48 h the read counts obtained on G_75_-L_25_, G_90_-L_10_ to those obtained on G_100_, or L_100_. In addition read counts obtained on L_100_ are also compared to those obtained on G_100_. This approach, sketched in the circuit design displayed in the Additional file 2, leads to ten possibilities for a gene to be identified as differentially expressed.

#### Gene expression matrix construction

For clustering and network inference, the establishment of a relevant gene expression matrix is needed.

For this purpose, we used results from the differential analysis. More precisely, we selected the subset of genes which are identified as differentially expressed in at least one on the ten studied comparisons. We decided to remove genes having at least one missing value over the ten comparisons. Doing this, we selected 650 genes for which a complete expression profile was available, composed of ten log_2_ expression ratios values leading to the gene expression matrix used to carry out the clustering. We note that, in this matrix, the fold change is computed on the average of the read counts across the biological replicates for a given condition (test or reference). For the network inference part, we choose to deal with a slightly modified version of this expression matrix, while keeping the same initial set of the 650 DE genes. To enforce the relevance of the metric used in network inference methods, we chose to deal with all biological replicates for the tested conditions while all reference conditions were pooled, with glucose or lactose pure are chosen as reference conditions. In other words, the log fold change is computed between the read count coming from a biological replicate of the test condition and the averaged read counts of the reference condition. Hence, for a given comparison, we obtained as many log fold changes as biological replicates. In order to harness the variability caused by this approach, we removed genes for which a biological replicate has a null read count. As a result, the final matrix contains 593 genes, where for each gene the expression profile contains 32 components. This procedure allows us to deal with expression profiles having a sufficient number of components to obtain a more reliable inferred network.

### Clustering and functional analysis of differentially expressed genes

#### Clustering

As previously mentioned, clustering is performed on the 650 genes. Each gene is characterized by its ten-component expression profile. The following approach was completely performed using the Multi Experiment Viewer (MeV) software [85]. Firstly, a hierarchical clustering allows us to estimate the optimal number *K* of clusters hidden in the data. By choosing the Euclidean distance metric and the average linkage method, results suggest *K* = 5 clusters. Then, the *K*-means algorithm (originating in [86]) is preferred in order to obtain a final gene classification. As this method is sensitive to initialization, we performed ten independent runs of *K*-means with random initialization; the Euclidean distance is used for each run. Results are subsequently aggregated into five consensus clusters. The aggregation is constrained by a co-occurrence threshold, fixed to 80%. As a result, the 650 genes are completely classified into five clusters and no unassigned cluster was found.

#### Functional analysis

A functional analysis was performed throughout a full expert annotation of the classified genes. For this purpose, each gene present in the clustering was manually curated using the *Trichoderma reesei* Gene Ontology (GO) annotation from the Joint Genome Institute (JGI) [87, 88], in terms of biological process and molecular function. Functions of genes, for which no process or nor function is found, are predicted by similarity to orthologous genes, when available in other fungal taxa. For this purpose, orthologs are determined *via* FungiPath and FungiDB. Note that by convention in this manuscript, we shall denote by *Tr* –XXX the gene in *T. reesei* for which the ortholog in an other specie is XXX. Otherwise, genes are labeled as unknown. This functional annotation allows us to manually provide meaning to clustering results.

### Network inference and promotor analysis

#### Network inference

Network inference was performed using the gene expression matrix containing 593 genes (and 32 differential expression levels) as input. We firstly obtained a complete weighted network 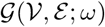, linking all genes 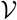 by links 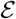 with weight *ω*. This step was performed thanks to the CLR (Context Likelihood of Relatedness) algorithm [61]. The weights *ω*_*i,j*_, affected to each pair (*i, j*) of genes, are based on the mutual information metric which quantifies the mutual dependence or the information shared between expression profiles of genes *i* and *j*. From this complete gene network, a threshold selects the most relevant gene links. For this purpose, we used the network enhancement algorithm BRANE Cut [40]. Briefly, each edge *e*_*i,j*_ in the complete network is labeled by a variable *x*_*i,j*_ set to 1 if the link has to be in the final network, and 0 otherwise. By optimizing a cost function over the variable 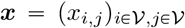, the minimizer ***x***\* gives us the optimal set of links on the final graph. In order to select the relevant links, biological and structural constraints are encoded in the cost function. Indeed, in addition to favoring strongly weighted edges, this post-processing method prefers links around labeled transcription factors. Moreover, thanks to an additional constraint, links between a gene and a couple of transcription factors, if this latter couple is identified as co-regulator, are also preferentially selected. As a result, we obtain an inferred network composed of 161 genes and 205 edges.

#### Promoter analysis

The promoter analysis was performed using the Regulatory Sequence Analysis Tools (RSAT) software [63]. From each set of genes to study (linked to a specific TF), promoter sequences from −1 to −1000 upstream bases are retrieved using the *retrieve sequence* tool. From these sequences, a detection of over-represented oligonucleotides was performed thanks to the *oligo-analysis* tool. We used the reference sequence set of *Trichoderma reesei* as background model. As mentioned in [89], this choice is driven by the fact that the input sequences (the query) are a subset of a larger collection (the reference). As a result, we obtain a list of over-expressed oligonucleotides (from hexa- to octo-) and several larger motifs assembled from the previous ones using the *pattern assembly* tool. Significance and count matrices are also obtained at this stage and lead to the establishment of sequence logo binding motifs. In order to detect the occurrences of the previously discovered patterns, we used the *string-based pattern matching* (dna-pattern) tool. It provides a list of features indicating the positions of the motifs in the input sequences. A suitable way to deal with this data is to visualize them using the *feature map* tool. From the feature map, the presence of overlapping close motifs is commonly a good indication for the relevance of the discovered motif. This methodology hints at supposing that the set of initial tested genes detains a binding site of the linked TF. From the given occurrences, we also computed the average number of discovered sites on the tested subset of genes. Then, in order to give a statistical significance, we performed two statistical analyses: one based on the promoter sequence of the whole genome, the other based on the a set of random promoter sequences. For both statistical analysis, the occurrences are also computed and averaged over the number of involved sequences. Then a *t*-test was carried out in order to deem significance (or not) to the average number of discovered sites. The significance is given for a *p*-value lower than 0.05.

## Acknowledgements

We would like to thank Dimitri Ivanoff, Sabine Prigent and Thiziri Aouam for technical assistance. This work was supported by the France Génomique national infrastructure, funded as part of the “Investissements d’Avenir” program managed by the Agence Nationale de la Recherche (contract ANR-10-INBS-09)

## Authors’ contributions

AP analyzed and interpreted the RNA-seq data through a series of bioinformatics analyses (DE, clustering, network inference, GO enrichment, promoter analyses) and also participated to the redaction of the article. LD collaborated to the design of the experiment and reviewed the manuscript. FBM coordinated the study, interpreted the data analyses and reviewed the manuscript. EJ and AM designed and supervised the study and drafted the manuscript. CB, CF and SP carried out the RNA-seq experiments and bioinformatics.

## Additional Files

Additional file 1 — Study of the biomass concentration during the *fed-batch*

This PNG file contains experimental results regarding the study of the Rut-C30 biomass concentration at 0 h, 24 h, 48 h and 120 h during the *fed-batch* on G_100_, G_75_-L_25_ , G_90_-L_10_ and L_100_ .

Additional file 2 — Circuit design

This PDF file contains an illustration of the methodology used to perform the differential analysis.

Additional file 3 — List of mutated and/or differentially expressed genes

This Excel file contains two sheets. In the first one, we found the list of differentially expressed genes and contains information regarding gene name, gene function, orthologs in various species (*S. cerevisiae*, *A. nidulans* and *N. crassa*), whether the gene is a transcription factor, expression ratios and the label of the cluster to which it belongs. In the second sheet, there is the list of mutated genes in Rut-C30, by comparison to QM6a, and the ones which are identified to be differentially expressed in our conditions.

Additional file 4 — Promoter analysis of *clr2*

This Excel file contains three sheets. The first one gathers results regarding the promoter analysis of *clr2* based on results obtained in the sub-network SubN1 generated by BRANE Cut [40]. The second sheet displays the pattern feature map while the third one contains the statistical analysis regarding the discovered promoter sequence.

